# Spatiotemporal AI modeling of gene regulation and variant effects in the developing human brain

**DOI:** 10.64898/2026.10.09.757610

**Authors:** Yun Hao, Christopher Y. Park, Chandra L. Theesfeld, Olga G. Troyanskaya

**Affiliations:** Center for Computational Biology, Flatiron Institute, New York, NY, USA; Lewis-Sigler Institute for Integrative Genomics, Princeton University, Princeton, NJ, USA; Princeton Precision Health, Princeton University, Princeton, NJ, USA; Department of Computer Science, Princeton University, Princeton, NJ, USA; Department of Molecular Biology, Princeton University, Princeton, NJ, USA

## Abstract

Understanding how genetic variation shapes gene regulation in the developing human brain remains a central challenge in neuroscience, particularly in transient cell types and developmental stages where data are limited. We present Seal, an interpretable AI transfer learning framework for genome-based modeling of gene expression and variant effects with spatiotemporal resolution across 26 brain regions, 30 cell types, and seven developmental stages. Seal predictions enable characterization of both germline and somatic brain variants, and provide mechanistic interpretation by linking sequence variants to transcriptional regulators in spatiotemporal context. Applying Seal to genome-wide association studies, we identify cell types and developmental windows relevant to neuropsychiatric disease risk, revealing shared and distinct regulatory architectures across six neuropsychiatric conditions that align with clinical trajectories. In the Simons Simplex Collection autism cohort whole genome data, Seal uncovers a significant burden of de novo regulatory variants in transient fetal excitatory neurons, revealing a cell-type and developmentally specific regulatory burden. By connecting genetic variation to spatiotemporal regulatory programs, Seal offers a general framework for uncovering mechanisms of human brain development and disease.

## Introduction

Unraveling the regulatory mechanisms of gene expression is a critical task in human genetics and precision medicine^1–4^. A major challenge is deciphering the spatiotemporal dynamics of gene expression and interpreting transcriptional effects of genome variation in specific contexts of cell states^5–7^. This is intractable to fully address experimentally due to both genomic diversity and the transient nature of many cell states involved in the spatiotemporal response, such as during development^8^, aging^9^ or infections^10^.

Addressing this challenge thus requires computational models capable of linking genomic sequences and variant effects to spatiotemporally resolved expression consequences. Current deep learning frameworks, such as ExPecto^11^, ExPectoSC^12^, Enformer^13^, Borzoi^14^, and others^15–18^ have shown significant promise in predicting tissue- and cell-type-specific gene expression and variant effects. However, they fall short of making predictions in highly specific contexts of cell states, largely due to inherent limitations in their architectures and external constraints related to data availability. Existing model architectures require substantial amounts of training data for each context, which is often experimentally impractical due to technical difficulty and lack of post-mortem samples. To address the challenge, we introduce a sequence-based framework with transfer learning architecture designed to be both robust and accurate for highly specific spatiotemporal contexts. Our framework is tailored to the specific contexts of brain development, addressing a critical need in the field, though the framework is general and can be expanded to other contexts.

Understanding the regulatory landscape of brain development is critical to deciphering the molecular architecture of neuropsychiatric conditions^19,20^. While genome-wide association studies (GWAS) have identified many candidate loci, their specific regulatory and phenotypic effects on brain development remain largely unknown ^21,22^. Human brain development relies on the precise execution of spatiotemporal gene expression programs across numerous, often transient, cell states from early fetal stages through to adult maturation^23–25^, for example, early fetal astrocytes or neonatal interneurons. Due to the complexity and diversity of neural cell states, representations for these specific states are rare even within comprehensive datasets like PsychENCODE^26^, BICCN^27^, and Brain Cell Atlas^28^. This poses a significant challenge for systematic sequence-to-expression modeling.

We address this challenge with Seal, a deep learning framework for predicting context-specific variant effects on expression *ab initio* from sequence. The Seal framework uses a transfer learning-based neural network architecture to leverage information from deep and numerous gene expression profiling for adult brain regions to inform fine tuning for contexts with very limited data, especially that from brain cell types early in development. Transfer learning, which has been successfully applied to other areas of biological research^29–31^, enables Seal to boost performance in sequence-based prediction of cell-state-specific gene expression, covering over 800 brain regions and cell type contexts.

The Seal framework can facilitate and accelerate the investigation of spatiotemporally resolved regulatory architecture underlying brain development and neuropsychiatric conditions. Along with the variant effect predictions, the Seal framework provides interpretation by linking predicted variant effects to important biochemical regulators such as transcription factors (TFs) and histone marks. We applied the Seal framework to characterize disease- and trait-associated regulatory disruptions in GWAS data, and uncovered the developmental timing and relevant cell types contributing to neuropsychiatric conditions. Moreover, applying Seal to whole genome sequencing data from the SSC autism cohort, we discovered that *de novo* variants in individuals with autism cause significant functional burden in transient cell states of fetal excitatory neurons. In summary, we provide an end-to-end computational framework for *in silico* prediction of context-specific variant effects in highly specific biological contexts, including in data-scarce settings.

## Results

### Seal is a transfer-learning-based, spatiotemporally-resolved sequence-to-expression framework

To enable sequence-based prediction of gene expression in the specific contexts of brain development, we built a deep learning approach that can train on gene expression data derived from both bulk RNA-seq profiling and single-cell RNA-seq profiling (Fig. 1a and Methods). Our new approach, Seal, adopts a transfer learning framework to enable accurate modeling of limited, highly context-specific gene expression data in 802 diverse contexts of brain development from early fetal stages through adulthood.

**Fig. 1.**
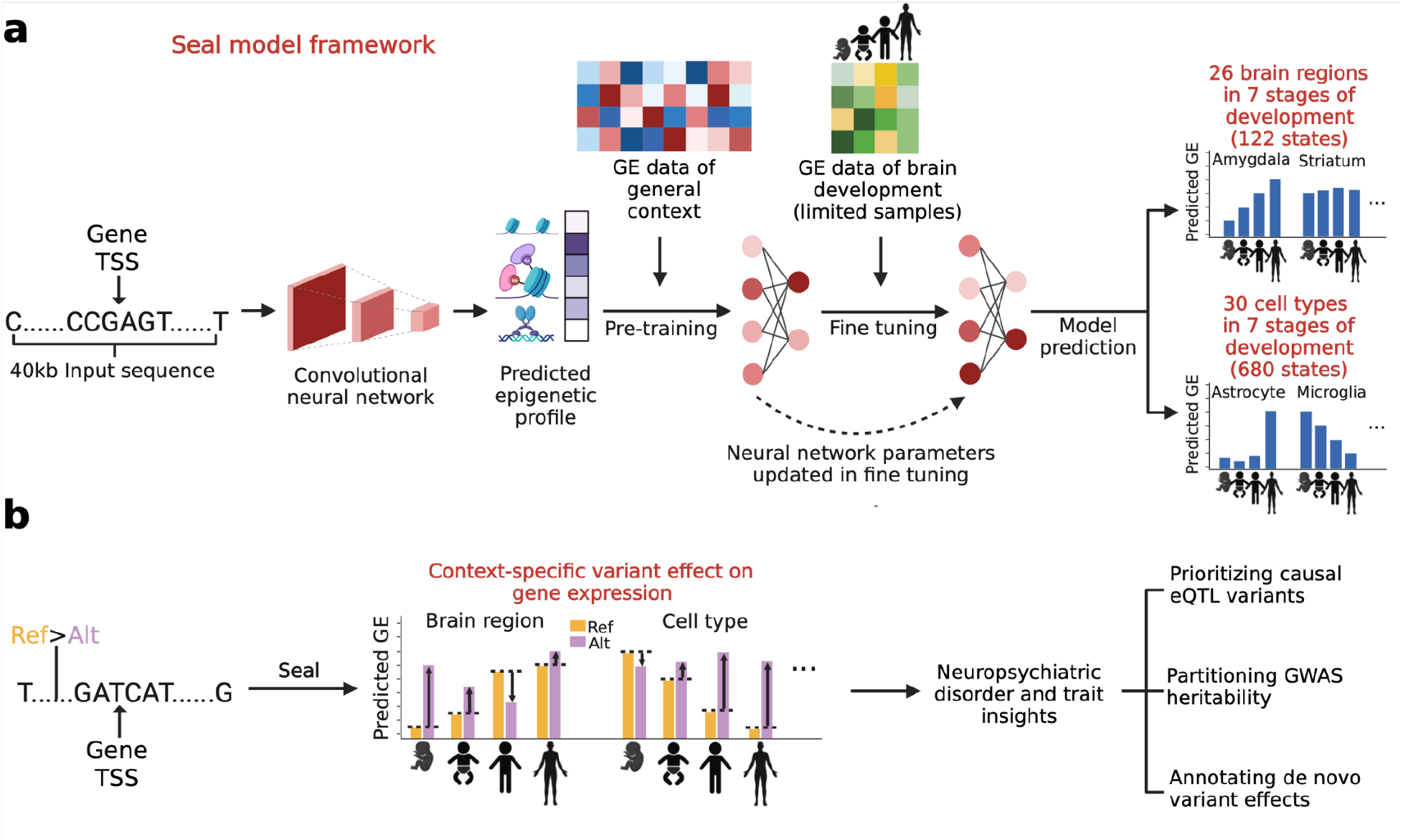
Sequence-based transfer learning model predicts variant effects on gene expression in 802 contexts of brain development. **a**, Overview of the Seal **S**equence-to-**e**xpression tr**a**nsfer **l**earning framework. Seal takes in 40kb sequence centered around the transcription start site (TSS) of each gene. The model contains three modules, a convolution neural network module that transforms input sequence to epigenetic features, a pre-training module that learns a multi-layer perceptron model for relating epigenetic features to gene expression values under general contexts, and lastly, a fine-tuning module that updates the parameters of pre-trained model for predicting gene expression of specific contexts: 802 brain-specific states in seven stages of development, including 122 states for 26 brain regions and 680 states for 30 brain cell types (evaluation detailed in Fig. 2). **b**, Application of Seal framework for predicting context-specific variant effects on gene expression. Variant effect in each context is quantified by the difference in predicted expression values of reference (ref) and alternative (alt) allele. The Seal framework can accomplish important tasks in human genetics: prioritizing causal expression quantitative trait loci variants (detailed in Fig. 3), understanding the regulatory architecture of brain-associated traits, such as partitioning GWAS heritability (detailed in Fig. 4), whole genome analysis for defining the functional burden of variants, such as annotating the cell-state-specific effects for rare de novo variants from disease cohort (detailed in Fig. 5).

The Seal framework first uses a convolutional neural network to generate an epigenomic representation of the 40kb sequence centered around the transcription start site (TSS) of each gene. The resulting *in silico* epigenomic profile embeds 2002 genome-wide biochemical regulators of gene expression, including DNA accessibility, histone marks, and transcription factor binding. Second, in a pre-training step, Seal learns latent space representations of epigenomic features by training a multi-layer perceptron neural network model with these 2002 epigenomic profile embeddings to predict gene expression for 31 brain regions and derived cell lines (Supplementary Table 1), leveraging the rich data generated by GTEx, ENCODE, and Roadmap projects, with an average of 124 samples per region. Third, Seal is fine-tuned with highly specific, but limited-availability gene expression profiling data for the seven stages of brain development (early fetal, mid fetal, late fetal, infancy, childhood, adolescence, adulthood; Supplementary Table 2), updating weight parameters of the latent space learned from pre-training.

Seal predictions are interpretable because predicted expression variant effects can be traced back to specific biochemical impacts on 2,002 epigenomic features. Seal provides feature attribution scores for biochemical interpretation on how a variant regulates gene expression in each cell state, through disruption of TF binding (690 features), modification of histone marks (978 features) or chromatin accessibility changes (334 features). This can be used to develop detailed hypotheses on regulatory changes by variants.

### Transfer learning is important for Seal’s predictive performance

The transfer learning framework is able to leverage information from the pre-training step to inform the fine-tuning step. The transfer learning framework significantly improved the overall model performance of Seal as compared to training with all data in one step (*P* = 0.01, Wilcoxon signed-rank test; Supplementary Fig. 1a; Methods). The improvement is especially high for genes involved in neurodevelopment processes (*P* = 1.1*10^−18^, Wilcoxon signed-rank test; Supplementary Fig. 1b). Expression of these genes is more challenging to predict due to high variability across different stages of development.

### Seal demonstrates high accuracy and specificity in sequence-based gene expression prediction

Seal has >8-fold higher coverage of brain-specific contexts (e.g. early fetal dorsolateral prefrontal cortex, neonatal interneuron) compared to state-of-the-art sequence-to-expression models (Fig. 2a), especially in the early stages of brain development. Evaluated on the test set comprising genes of chromosomes 8 and 9 (held-out at all stages of training), Seal achieved an average Spearman correlation of 0.808±0.002 between predicted and observed expression across all 802 contexts, and an average area under the receiver-operating characteristic curve (AUROC) of 0.987±0.001 (Fig. 2b and Methods), demonstrating that Seal can accurately predict the context-specific gene expression levels from sequences.

**Fig. 2.**
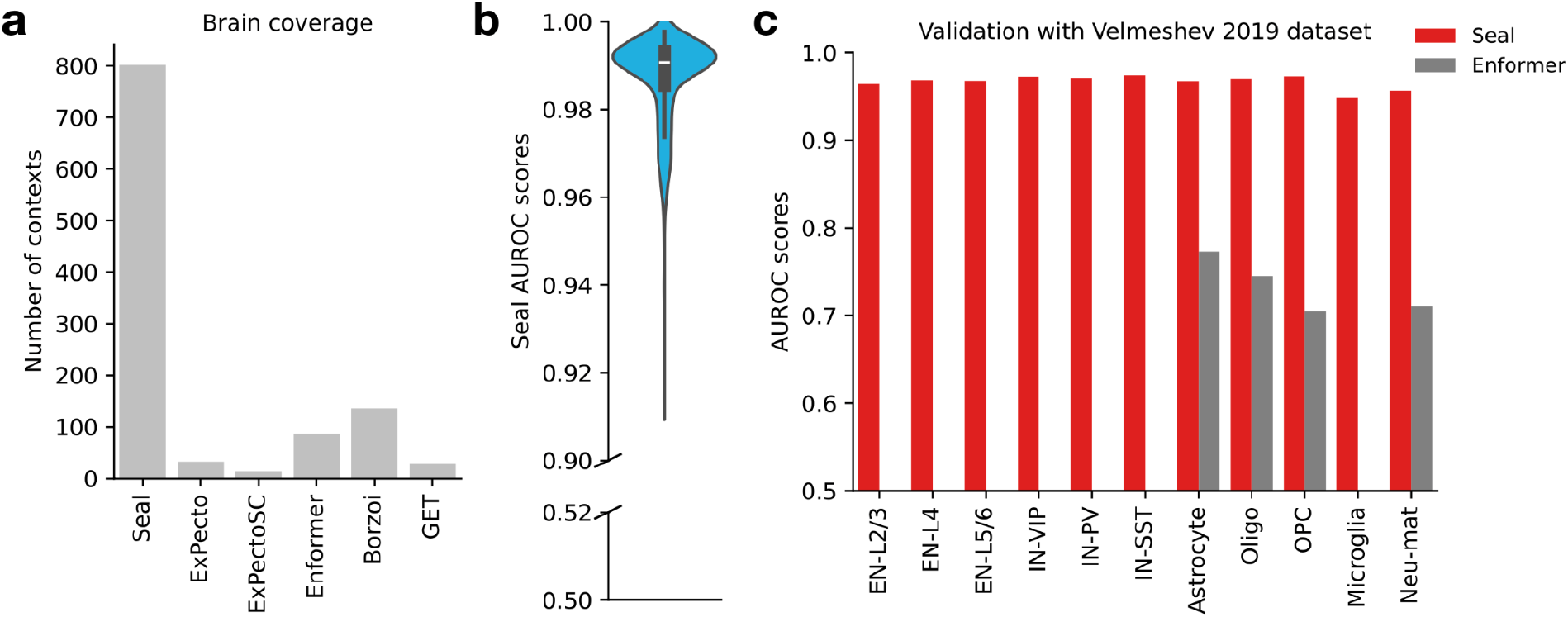
Seal provides accurate sequence-based expression predictions for 802 brain-specific states. **a**. Seal has a much broader coverage of brain-specific contexts compared to five state-of-the-art sequence-to-expression models. The bar shows the number of brain-specific contexts covered by each model. ExPecto and ExPectoSC are non-transfer-learning-based frameworks previously developed by our group. Enformer and Borzoi are self-attention-based models with Transformer-like architecture. GET is a DNA foundation mode for transcriptional regulation. **b**, The sequence-based gene expression predictions by Seal are highly correlated with observations. For each specific context, an AUROC score for predicting highly and lowly expressed genes was computed for the 1,912 genes on holdout chromosomes 8 and 9. The violin plot shows the distribution of AUROC scores across all 802 contexts. **c**, Seal outperforms Enformer in predicting context-specific gene expression of brain development. Sequence-based predictions by Seal and Enformer were benchmarked against two independent single-cell RNA-seq datasets of the fetal brain (Validation with the Velmeshev 2019 dataset is shown here, Validation with the Bhaduri dataset is shown in Supplementary Fig.2). The bar plot shows the computed AUROC of each identified cell state. Seal can make predictions for all the identified cell states while Enformer cannot. Cell states where Enformer cannot make predictions were left blank (no grey bar). (abbreviations, EN: excitatory neuron; IN: interneuron; Oligo: oligodendrocyte; OPC: oligodendrocyte precursor cell; Neu-mat: maturing neuron)

We then benchmarked Seal against two external scRNA-seq datasets of fetal brain: Velmeshev *et al*^*32*^ and Bhaduri *et al*^*33*^ (Methods). Without training on these data, Seal exhibits consistently high accuracy for gene expression prediction across all cell states (Fig. 2c and Supplementary Fig. 2), significantly outperforming Enformer^13^ on the contexts for which Enformer also provides coverage. Specifically, Seal achieved an average AUROC of 0.967±0.003 for the Velmeshev *et al* dataset (vs. 0.733±0.026 for Enformer), and 0.940±0.006 across all cell types from the Bhaduri *et al* dataset (vs. 0.747±0.019 for Enformer). Importantly, Seal has a much broader coverage of brain-specific cell states compared to Enformer (e.g. fetal microglia, EN-L2/3, EN-L4, EN-L5/6, IN-VIP, IN-PV, IN-SST; Fig. 2c), all of which are missing from Enformer predictions.

### Seal predictions prioritize expression-altering eQTL variants for brain development

Seal is not biased towards variants with high allele frequency and thus can probe the expression effect for any type of variants *ab initio*: common, rare, or previously unobserved in sequencing studies. The *ab initio* nature of sequence-based prediction by Seal makes it unconfounded by linkage disequilibrium (LD), making it complementary to quantitative trait loci (QTL) methods. Therefore, Seal can infer casual variant effects on gene expression, addressing another major limitation of prior quantitative genetics approaches.

To evaluate the performance of Seal on variant effect prediction, we probed the expression effects of 743,810 brain-associated eQTL variants from two sources, including 80,594 eQTLs in three fetal stages of prefrontal cortex discovered by Wen *et al*^*34*^ and 663,216 eQTLs in seven adult brain regions discovered by the GTEx consortium^7^. Facilitated by the broad coverage of brain-specific contexts, Seal extends the eQTL variant effect prediction to more specific brain regions and stages of development. For example, while identification of eQTLs in Wen et *al*^*34*^ was limited by sample availability to prefrontal cortex (PFC) in three fetal stages, Seal extends the number of contexts that effects can be predicted for to 12 different fetal PFC contexts: including four PFC regions (medial, orbital, dorsolateral, and ventrolateral) for each of the three stages of fetal development (early fetal, mid fetal, and late fetal). The predictions of Seal include the direction of expression change (up-regulated and down-regulated) and effect size that can be used for variant prioritization. Since the vast majority of eQTLs have no effects on gene expression (<1% of all GTEx eQTL variants were estimated to be causal^35^), we then examined the extent to which Seal can complement quantitative genetics approaches by prioritizing the true causal eQTLs (Fig. 3a and Methods). Seal correctly predicted the direction of expression change for top prioritized eQTL variants, yielding consistently high accuracy scores across three evaluated stages (an average accuracy of 0.82, Fig. 3b-d) and seven brain regions (an average accuracy of 0.84, Supplementary Fig. 3).

**Fig. 3.**
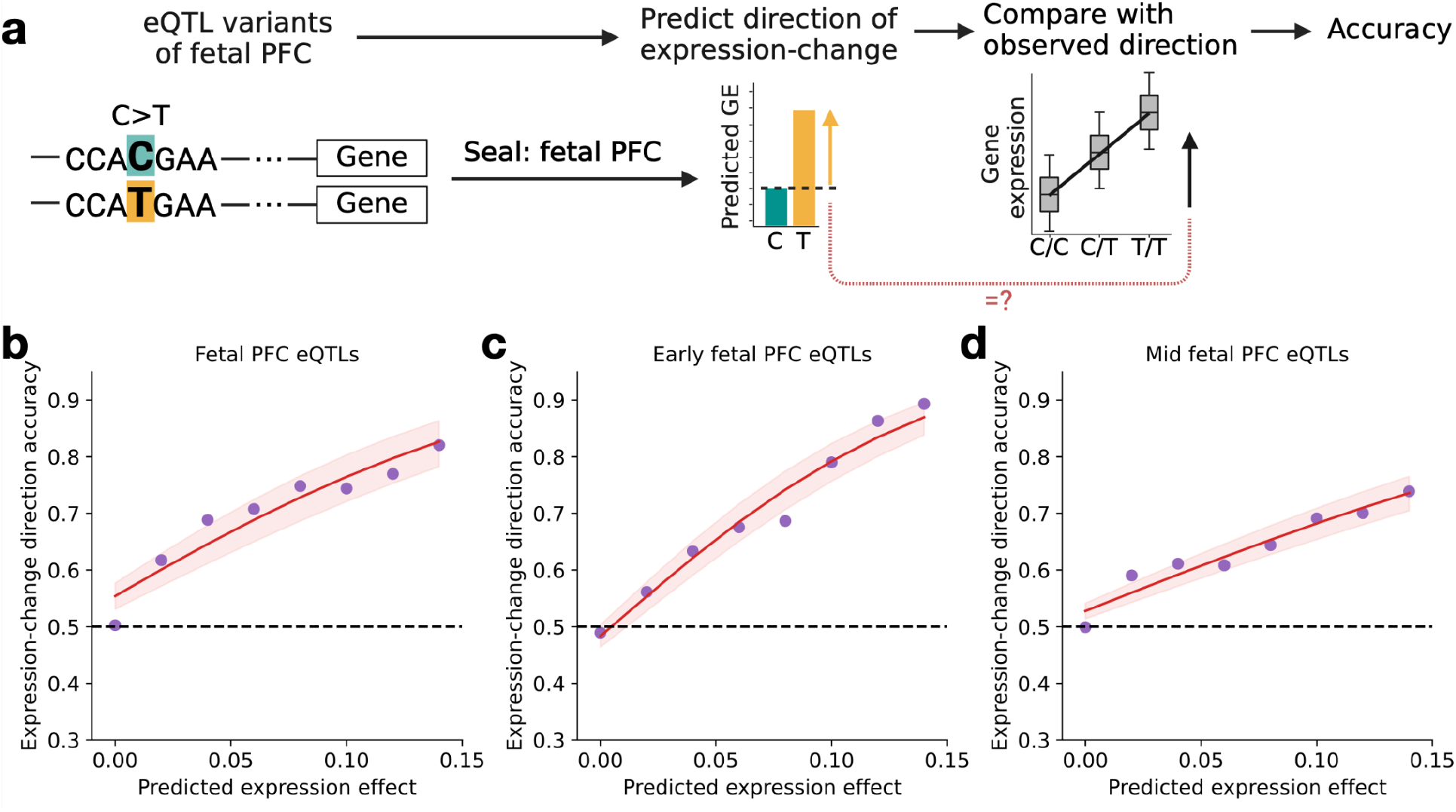
Seal prioritizes expression-altering eQTL variants for brain development. **a**, Workflow of analyzing expression quantitative trait loci (eQTL) variants of brain development with Seal framework. The Seal prediction for a specific eQTL variant includes both the direction of expression change (up-regulated and down-regulated), as well as the effect size which can be used for variant prioritization. Here, Seal was applied to predict the expression-change direction of eQTL variants in the specific context where the eQTLs were identified. Three sets of eQTL variants were analyzed using this workflow: fetal prefrontal cortex (PFC) eQTLs (shown in **b**), early-fetal PFC eQTLs (shown in **c**), and mid-fetal PFC eQTLs (shown in **d**). Results for the seven sets of eQTL variants discovered from adult brain regions can be found in Supplementary Fig. 3. The accuracy score was computed by comparing the predicted direction with the observed direction of each eQTL variant. **b-d**, Expression-change direction accuracy increases with the predicted magnitude of variant effect, suggesting eQTLs with strong Seal-predicted effects are more likely to be causal. Each dot in a plot represents the computed accuracy (y axis) for variants above a predicted effect magnitude threshold (x axis, measured by the absolute value of log fold change). Each curve was fitted using the computed accuracy scores at eight different thresholds, with the 95% confidence interval represented by the shaded area. Seal correctly predicted the direction of expression change for top prioritized eQTL variants with high accuracy across all three stages.

### Seal recapitulates temporal-specific regulatory programs of brain structural traits

Seal’s broad spatiotemporal context coverage and accurate prioritization of eQTL variants suggest Seal predictions can be used to study the genetics of brain traits. To test this hypothesis, we used Seal variant effect predictions as a set of genome-wide annotations to partition heritability in well-powered GWAS studies using LD score regression (LDSR)^36^, which can link brain traits to potential brain region state. LDSR is a statistical framework for partitioning heritability into various functional annotations, while directly modeling the linkage disequilibrium (LD) between variants^36^. We estimated the enrichment of heritability explained by variants perturbing gene expression in each brain region state for two GWAS of brain structural traits^37^: cortical mean thickness and cortical surface area (Methods). The sequence-based variant annotations by Seal included 122 spatiotemporal contexts that involve 26 brain regions and seven stages of development (Supplementary Table 3).

We found differential enrichment of fetal and adult spatiotemporal contexts for these two traits. Notably, we found temporal context differences in genetic contribution: cortical mean thickness showed higher enrichment of heritability among variants perturbing gene expression in the adult stage (*P* = 6.3*10^−10^, Mann-Whitney U test; Fig. 4a), whereas cortical surface area showed higher enrichment of heritability among variants perturbing gene expression in the fetal stage (*P* = 5.5*10^−4^, Mann-Whitney U test; Fig. 4a). This is consistent with previously published work that showed active adult-specific regulatory elements were enriched for cortical mean thickness and fetal-specific elements were enriched for cortical surface area^37^. We also found spatial differential enrichment among brain regions for cortical surface area, with higher enrichment of heritability among variants perturbing gene expression in transient fetal regions (e.g. ganglionic eminence) compared to fetal brain regions that persist until adulthood (*P* = 5.5*10^−5^, Mann-Whitney U test; Fig. 4b). This finding further highlights the strong influence of fetal-specific transcriptional regulation on cortical surface area. The detailed sequence-based annotations by Seal combined with LDSC analysis can provide fine resolution for the causal contribution of specific regions and cell states at distinct developmental times.

**Fig. 4.**
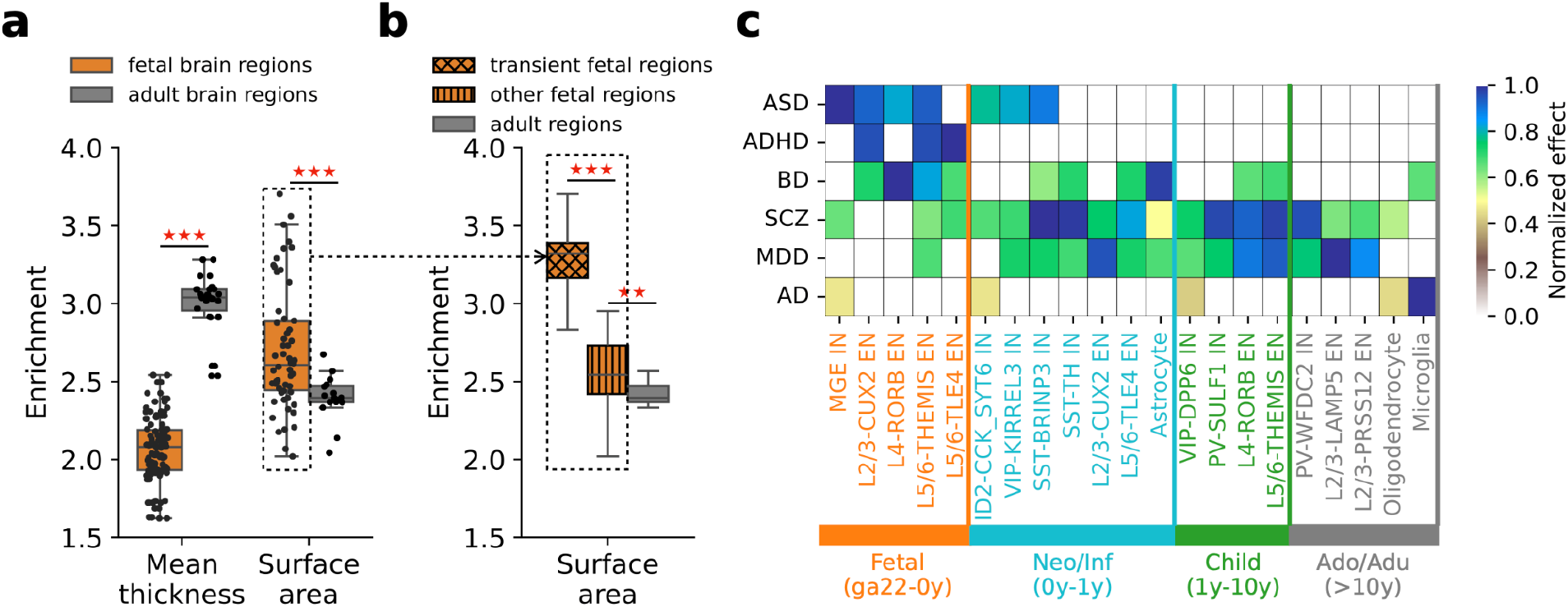
Seal-based partitioning of GWAS heritability reveals the developmental timing and important cell types that contribute to brain-associated traits. **a-b**, Seal-based partitioning recapitulates temporal-specific regulatory programs of brain structural traits. LD score regression (LDSR) was implemented to estimate the proportion of heritability explained by each brain-region state for the two traits. The effect sizes (estimated enrichment, y-axis) of brain regions were compared between fetal stage and adult stage for two brain structural straits (shown in **a**): cortical mean thickness and cortical surface area (x-axis). Higher enrichment of heritability was observed within fetal brain regions for cortical surface area (P = 5.5*10^−4^, Mann-Whitney U test), whereas the opposite trend was observed for cortical mean thickness (P = 6.3*10^−10^, Mann-Whitney U test). Further analysis (shown in **b**) highlights the strong influence of fetal-specific transcriptional regulation on cortical surface area. Fetal brain regions were further split into transient fetal regions and other regions that persist into later stages of development. Higher enrichment was observed in transient fetal compared to other regions (P = 5.5*10^−5^, Mann-Whitney U test). **c**, Seal-based partitioning reveals both shared and distinct regulatory architecture among neuropsychiatric conditions. Following the same procedure, LD score regression (LDSR) was implemented to estimate the proportion of heritability explained by each cell state for six neuropsychiatric conditions (x-axis, ASD: autism spectrum disorder, ADHD: attention-deficit hyperactivity disorder, BD bipolar disorder, SCZ: Schizophrenia, MDD: major depressive disorder, AD: Alzheimer’s disease). For each disorder, the estimated effect sizes were normalized into scores between 0 and 1 (scale indicated by the color of each cell) and the top five most enriched cell states (by FDR) were selected for this heatmap (21 in total due to overlaps). The selected cell states were sorted and colored by stages of development (y-axis, Neo/Inf: neonatal and infancy, Ado/Adu: adolescence and adulthood, ga: gestational age). The heatmap shows the significantly enriched cell states (FDR < 0.05) for each condition, with the color scale proportional to the estimated effect size. (EN: excitatory neuron; IN: interneuron; MGE: medial ganglionic eminence; ID2: inhibitor of DNA binding 2; VIP: vasoactive intestinal peptide; SST: somastatin; PV: parvalbumin)

### Seal identifies stages and cell types linked to risk of neuropsychiatric conditions

Neuropsychiatric conditions involve the disruption of transcriptional regulation in specific cell types and stages of brain development^20^. Seal can systematically link regulatory variants identified in GWAS studies to their effects in cell states of brain development, predicting impacted cell types at specific stages of development. We applied LDSR^36^ to estimate the enrichment of heritability explained by variants perturbing gene expression in each SEAL-modeled cell state for six conditions: autism spectrum disorder (ASD), attention-deficit/hyperactivity disorder (ADHD), Bipolar disorder (BD), Schizophrenia (SCZ), Major depressive disorder (MDD), and Alzheimer’s disease (AD) (Methods). The sequence-based variant annotations by Seal covered 82 spatiotemporal contexts that involve 18 cell types of prefrontal cortex and six stages of development from mid-late fetal to adulthood (Supplementary Table 4).

This analysis revealed aspects of regulatory architectures that are both shared and distinct among these conditions (Fig. 4c). Notably, the two neurodevelopmental conditions, ASD and ADHD, only showed significant enrichment of heritability at fetal, neonatal, and infancy stages. By contrast, other conditions (BD, SCZ, MDD, and AD) showed impact on at least some cell types across all six developmental stages, with higher enrichment observed within childhood, adolescence, and adulthood. The result is consistent with the onset and timing of these conditions, with ASD and ADHD exhibiting much earlier age of onset (early to mid childhood) compared to other conditions (late adolescence to adulthood)^38^. This analysis also recapitulated brain cell types that are known to be involved in the studied neuropsychiatric conditions (Figure 4c). ASD showed significant enrichment of heritability among variants perturbing gene expression in medial ganglionic eminence (MGE IN, FDR = 0.016), deep layer 5 and 6 excitatory neurons (L5/6-THEMIS EN, FDR = 0.024), and superficial layer 2 and 3 excitatory neurons (L2/3-CUX2, FDR = 0.034), all of which have prior evidence of association with ASD^39–41^. In contrast, AD was associated with microglia (FDR = 0.013) and oligodendrocytes (FDR = 0.022), consistent with extensive evidence suggesting the two glia cell types are morphologically and functionally impaired in animal models of AD^42–45^. Similarly, our analysis recapitulated connection of L5/6-THEMIS EN^46^ for MDD^46^ (FDR = 3.1*10^−3^), SST interneurons (SST-BRINP3 IN and SST-TH IN, FDR = 1.9*10^−4^ and 2.1*10^−4^, respectively) and Parvalbumin interneurons^47^ (PV-WFDC2 and PV-SULF1, FDR = 2.4*10^−4^ and 2.5*10^−4^, respectively) with SCZ^47^.

In addition to recapitulating existing knowledge, our analysis also identified potential novel disease-associated cell states, such as KIRREL3-expressing VIP interneurons (VIP-KIRREL3, FDR = 0.037) and BRINP3-expressing SST interneurons (SST-BRINP3 IN, FDR = 0.023) for ASD. While the two genes and cell types show relevance to neurodevelopment^48,49^, these precise cell states have not been linked to ASD. Such predictions provided by Seal enables development of new spatiotemporal-specific hypotheses for studying the mechanisms of neuropsychiatric conditions.

### ASD regulatory mutations show strong functional burden in fetal excitatory neurons

Seal enables analysis of whole genome sequencing data for cohorts and can generate mechanistic hypotheses for genetic dysregulation of specific cell states at precise developmental times. We used Seal to analyze regulatory variants from whole genomes to study gene expression regulation in ASD. Previously, we demonstrated the functional burden of *de novo* proband-specific mutations that perturb transcriptional and post-transcriptional regulation in the Simons Simplex Collection cohort^50^. Facilitated by the broad coverage of brain-specific contexts, Seal can further characterize the functional burden of *de novo* mutations in a cell-type- and state-specific manner (Fig. 5a; Methods). In total, we predicted the effects of 27,926 TSS-proximal *de novo* mutations from 1,781 ASD probands and 1,735 unaffected siblings in 82 cell states of the prefrontal cortex, involving 18 cell types and six stages of development from mid-late fetal to adulthood. Over all cell states, we observed higher predicted expression effects for mutations near known ASD risk genes, with significantly increased effect size for highest-confidence ASD risk genes^49^ (Supplementary Fig. 4).

**Fig. 5.**
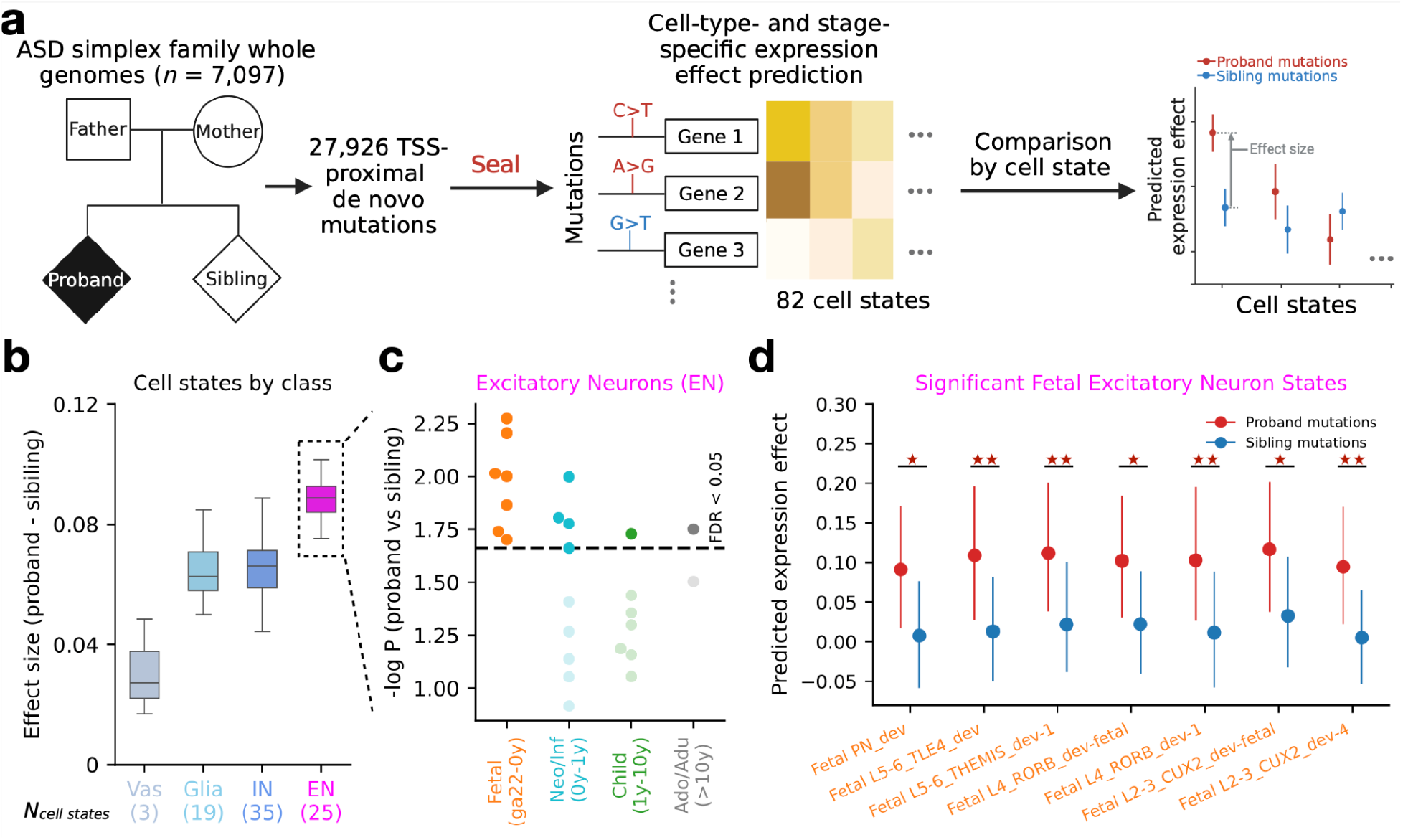
Cell-state-specific analysis of ASD regulatory mutations reveals strong functional burdens in fetal excitatory neurons. **a**, Workflow of cell-state-specific functional burden analysis with Seal framework. Whole genomes of 7,097 individuals from 1,790 autism spectrum disorder (ASD) simplex families were sequenced to identify de novo mutations in the ASD probands and unaffected siblings (27,926 mutations within 20kb of gene transcriptional start sites). Seal was applied to predict the cell-state-specific mutation effects on expression for 82 cell states of the prefrontal cortex. In each cell state, the predicted effects of proband mutations were normalized and compared against those of sibling mutations (one-sided Mann-Whitney U test). Effect size was computed as the difference in average predicted effects between proband and sibling mutations. The functional burden analysis result for all mutations is detailed in Supplementary Fig. 4. Results for a subset of mutations near ASD risk genes (SFARI category 1 and 2 genes, implicated in ASD by the presence of recurrent likely-gene-disrupting mutations) are shown in **b-d**. **b**, ASD proband mutations exhibit the highest effect size in cell states of excitatory neurons. All 82 cell states were grouped into four major cell classes (Vas: vasculature cells, Glia: glial cells, IN: interneurons, EN: excitatory neurons). The number of cell states within each class is shown in the parenthesis at the bottom. The boxplot shows the distribution of effect size (y-axis) among each major cell class (x-axis). **c**, ASD proband mutations exhibit strong functional burden in cell states of fetal excitatory neurons. The 25 cell states of excitatory neurons (class with the highest effect size in **b**) were further split, sorted, and colored by stages of development (x-axis, ga: gestational age, Neo/Inf: neonatal and infancy, child: childhood, Ado/Adu: adolescence and adulthood). The scatter plot shows the significance of functional burden (y-axis, negative log P-value of proband vs. sibling comparison) by stages of the cell state. The gray dashed line shows the significance threshold (FDR < 0.05). **d**, Comparison results for the significant fetal excitatory neuron states. The plot shows the comparison of Seal-predicted expression effects (y-axis) between proband (red) and sibling mutations (blue) for seven fetal excitatory neuron states (x-axis) with FDR < 0.05 in **c**. Each point shows the average predicted effects among mutations while the lines represent the 95% confidence interval. Asterisks denote the level of significance (one asterisk for 0.01 < P < 0.05, two asterisks for P < 0.01).

We then conducted cell-state-specific functional burden analysis for the 1,704 mutations near ASD-associated genes (SFARIgene category 1 and 2), comparing proband-specific mutations versus sibling-specific mutations (Supplementary Table 5; Methods). We found the strongest proband-specific burden in Excitatory neurons (EN) (Fig. 5b), with all EN cell types significantly affected at the fetal stage and with decreasing proportion affected through neonatal and infancy stages (Fig. 5c). Among the 13 significant excitatory neuron states, seven of them belong to the fetal stage, including L2/3-CUX2, L5/6-TLE4, and L5/6-THEMIS (Fig. 5d), all of which has been implicated in ASD by previous studies^40,41^.

Seal also provides mechanistic interpretation for its prediction, including how proband-specific mutations regulate gene expression via disrupting TF binding. For each of the 690 TF-binding-related epigenomic features in Seal, we computed the average attribution score across all 14,440 ASD-proband-specific mutations for the 37 cell states where proband-specific mutations exhibit significant functional burden of TF binding changes (FDR < 0.05). Notably, many well-known ASD-associated TFs were ranked among the top by feature attribution (Supplementary Table 7), including SFARI category 1 gene p300, YY, CTCF, FOXP2 and SFARI category 2 gene JARID1A. Other top-ranked TFs, such as GATA3 and ZNF263, are involved in neural maturation and brain development^51^, with potential implication in ASD^52^. Our findings suggest the hypothesis that ASD development in fetal excitatory neurons is at least partly due to variants that alter gene expression through disruption of ASD-associated TF binding.

## Discussion

We present Seal, a sequence-based transfer learning framework for prediction of variant effects on gene expression for 802 brain-specific states, including early stages of development with extremely limited training data. Seal provides expansive coverage of spatiotemporal cell states and interpretability of predictions, enabling biomedical scientists to study regulatory architecture and genetic basis of brain development and disease at unparalleled resolution.

We demonstrate the power of Seal in analysis of genetic variants from both GWAS and WGS studies, identifying excitatory neurons regulatory burden that is strongest at early fetal stages. It is tempting to speculate that these results propose specific cell states and stage (fetal L5/6 ENs) for initial dysregulation in excitatory neurons in the context of ASD.

The Seal transfer learning framework can be adapted for predicting variant effects in contexts of other tissues, stages, or species beyond the developing brain. For instance, the framework can be fine-tuned with the newly released scRNA-seq profiling of human aging brains^53^, facilitating the study of neurodegenerative conditions. Our study could be a blueprint for leveraging insufficient context-specific data to decipher the noncoding genome and understand its contribution in complex diseases.

## Methods

### Seal framework architecture

The training of Seal sequence-to-expression transfer learning framework consists of three sequential modules. First, a deep convolutional neural network (CNN) module transforms input sequences into epigenomic features. Then, a pre-training module learns a multi-layer perceptron (MLP) neural network model for relating epigenomic features to gene expression values within general contexts. Lastly, a fine-tuning module updates parameters of the pre-trained MLP model to predict gene expression values in specific cell state contexts.

The first training module of Seal employs the BELUGA CNN model^11^ to transform input sequences into epigenomic features. Specifically, Seal uses a sliding window with 200-bp step size to scan the 40kb sequence (+20kb upstream and −20kb downstream) centered around the representative TSS (defined by CAGE peaks^54^) of Pol II-transcribed genes. At each step, the BELUGA model takes in the 2000-bp sequence (200-bp sliding window plus the centered 1,800-bp surrounding context sequence) and outputs the predicted probabilities of 2,002 epigenomic profiles (978 histone mark, 690 TF-binding, and 334 DNase hypersensitivity profiles). In total, the scan of TSS-proximal sequence yields 200 spatial bins and 400,400 resulting features. Seal then employs exponential decay functions to spatially transform the predicted epigenomic features and reduce the dimensionality of feature space. For each epigenomic profile, the predictions from 100 spatial bins downstream of TSS (from −20kb downstream to TSS) are reduced to five features using 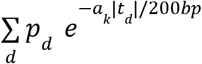 (*a*_*k*_ = 0.01, 0.02, 0.05, 0.1, 0.2) where *p*_*d*_ denotes the predicted probability at spatial bin *d* and *t*_*d*_ denotes the distance between the center of spatial bin *d* to TSS. Similarly, the predictions from 100 spatial bins upstream of TSS (from TSS to 20kb upstream) are reduced to five features using the same procedure. The use of exponential decay functions ensures that our transformation assigns higher weight to features near TSS. In total, the spatial transformation reduces the number of features 20-fold to 20,020.

The second training module of Seal pre-trains the sequence-transformed epigenomic features with a multilayer perceptron (MLP) neural network model for predicting gene expression values within general contexts (GTEX, ENCODE and ROADMAP, see below for details). The MLP model consists of three layers: an input layer of 20,020 nodes (taking in the 20,020 features derived from first module), a hidden layer of *N*_*h*_ nodes, and an output layer of *N*_*c*_ nodes (*N*_*c*_ denotes the number of contexts of interest). Specifically, the MLP model can be written as *Y = W*_*o*_*ReLu*(*W*_*h*_*X* + *b*_*h*_) + *b*_*o*_, where *X* denotes the input features, *W*_*h*_ and *b*_*h*_ denote the weight and bias matrices for the hidden layers, *W*_*o*_ and *b*_*o*_ denote the weight and bias matrices for the output layer. The MLP model is pre-trained with the gene expression data of general contexts, thus updating the *W*_*h*_, *b*_*h*_, *W*_*o*_, and *b*_*o*_ values. The pre-training process is equivalent to learning a latent space (i.e. the hidden layer) of the core epigenomic features that are responsible for driving transcriptional regulatory programs that are likely to be shared across cellular contexts. Since the number of latent features is far smaller than the number of input features, pre-training further reduces the dimensionality of feature space, as well as the number of model parameters to be updated in the next step.

The third training module of Seal fine-tunes the pre-trained model towards predicting gene expression values under specific contexts *Y*_*sc*_. Specifically, the hidden layer weight and bias matrices (*W*_*h*_ and *b*_*h*_) of the pre-trained MLP model are frozen after pre-training. Only *W*_*o*_ and *b*_*o*_ values are updated in this step. The fine-tuning process is equivalent to fitting context-specific regularized linear regression models against the latent features learned from pre-training. The simplified model can reduce the likelihood of overfitting on sample-limited expression data of each specific context. The predicted expression matrix *Y*_*sc*_ is evaluated against the observed expression matrix 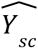 in a weighted mean squared error-based loss function 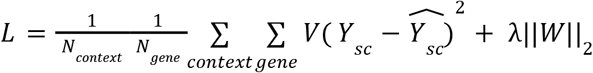, where λ denotes the L2 regularization coefficient and *V* denotes the gene weight matrix defined as

*ν* _*g*_*=s* (if the expression of *g* shows high variation across contexts),

*ν*_*g*_ *=* 1/*s* (if the expression of *g* shows low variation across contexts),

*ν*_*g*_*=*1(otherwise), and

*s* is a model hyperparameter. High variation is defined as above 70^th^ percentile while low variation is defined as below 10^th^ percentile. The rationale behind our gene-weighting strategy is to reduce the impact of ubiquitously expressed genes and ensure that Seal can capture the transcriptional regulatory programs in specific contexts.

### Context-specific prediction of variant effect on expression with Seal framework

The prediction of variant effects by Seal implements the first and third module sequentially while skipping the second module. The variant expression effect score is computed as the difference in the predicted expression levels between alternative and reference alleles (predicted log fold change). It can be written as 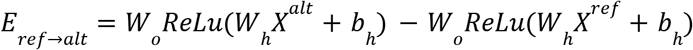, where *X*^*alt*^ and *X*^*ref*^ are the spatially-transformed epigenomic profile predictions of alternative allele and reference allele, respectively.

### Implementing Seal to predict spatiotemporal context-specific expression of brain development from sequence

We applied the Seal framework to train models predicting gene expression across brain development at both brain region and cell type resolution. For the pre-training step (second training module), we used the bulk RNA-seq expression profiling data of 31 brain regions and derived cell lines from GTEx^35^, ENCODE^55,56^, and Roadmap^57^(Supplementary Table 1). These brain regions were extensively profiled, with an average of 124 samples per region. For the fine-tuning step (third training module), we first concatenated three RNA-seq datasets (PsychENCODE^58^, Braun *et al*^*59*^, and Herring *et al*^*60*^) to compile a compendium of profiles that yield brain-region-state-specific and cell-state-specific gene expression data. The PsychENCODE^58^ bulk RNA-seq dataset (downloaded from http://development.psychencode.org) consists of 607 expression profiles of 26 brain regions from 41 individuals. Each profile contains the normalized RPKM values of gene expression. Based on the standard set by Kang *et al*^*61*^, the profiles of each region were grouped into seven developmental stages: early fetal (8 PCW-13PCW; PCW: post-conceptional weeks), mid fetal (13PCW-24PCW), late fetal (24PCW-38PCW), infancy (birth-1Y), childhood (1Y-12Y), adolescence (12Y-20Y), and adulthood (20Y-40Y), resulting in 122 stage-specific brain-region states. The gene RPKM values were averaged over the expression profiles of each state. The averaged RPKM values were then log-transformed and used as the input context-specific expression data 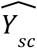 of the MLP model. The Braun *et al* dataset^59^ contains scRNA-seq profiling of whole brain samples in early fetal stage (5PCW-14PCW). In total, 616 cell states were identified in 12 brain cell types based on cell clustering. The Herring *et al* dataset^60^ contains scRNA-seq profiling of prefrontal cortex samples in six stages of development (defined by the authors), including mid to late fetal (22GW-birth, GW: gestational weeks), neonatal (birth-2M), infancy (2M-1Y), childhood (1Y-10Y), adolescence(10Y-20Y), and adulthood (20Y-40Y). In total, 86 cell states were identified in 18 brain cell types based on cell clustering. We concatenated cell states identified from the two studies and further removed states with fewer than 100 cells, resulting in 680 cell states of 30 cell types (598 states of early fetal, 82 states from mid-late fetal to adulthood; Supplementary Table 2). The gene CPM values were averaged over the cells of each cell state. The averaged CPM values were then log-transformed and used as the input context-specific expression data 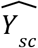 of the MLP model.

Genes on chromosomes 8 and 9 are held out as the test set in all stages of model training. The remaining genes are further split into a training set and validation set with a ratio of 7:1. This ensures that an independent validation set can be used for overfitting assessment. Each epoch of the training process consists of three steps. First, the training set (in mini-batches of size 32) is used for deriving gradients of network parameters via forward and backward propagation. The parameters are then optimized by Adam algorithm with learning rate *r*. Lastly, the validation set is used for determining whether the stopping criterion has been met (i.e. the loss function of the test set has not decreased for 20 epochs). In total, the Seal model has six hyperparameters: number of nodes in the hidden layer (*N* _*h*_), L2 regularization coefficient of pre-training (λ_*p*_), L2 regularization coefficient of fine-tuning (λ_*f*_), gene-weighting factor (*s*), learning rate of pre-training (r_*p*_), learning rate of fine-tuning (r_*f*_). We conducted a grid search among 2,430 hyperparameter combinations, then identified the optimal setting based on the mean square error of the validation set (supplementary table 7).

Using the held-out test set, we evaluated the gene-wise expression prediction performance of Seal. Specifically, we computed metrics on how well Seal separated highly expressed genes from lowly expressed genes in each specific context. We defined highly and lowly expressed genes as those with observed expression values in the top and bottom 25^th^ percentile, respectively. We labeled highly expressed genes as 1 and lowly expressed genes as 0. We then computed AUROC scores for each specific context by comparing the predicted expression percentile to ground truth.

We also benchmarked the gene-wise expression prediction performance of Seal on two external scRNA-seq datasets of fetal brain: Velmeshev *et al* ^32^ and Bhaduri *et al*^*33*^. We manually mapped cell states identified from the two datasets to those of Seal predictions by name. We computed gene expression values for each mapped cell state following the same procedure as described above. We utilized Seal and Enformer^13^ to generate sequence-based gene expression predictions for all mapped cell states. Enformer was implemented using code from the enformer-pytorch github repository (https://github.com/lucidrains/enformer-pytorch). For each cell state, we obtained gene expression predictions for the matched Enformer track by summing up predicted CAGE read counts at the TSS location of each gene (the same procedure described in the Enformer paper).

### Incorporating interpretation method in Seal framework to compute feature attribution scores for predicted variant effects

We computed variant effect attribution scores for the 20,020 input features of fine-tuned MLP model (third module of Seal) using the DeepLIFT method^62^. For each variant, input features for the reference allele were considered as “baseline instance” while input features for the alternative allele were considered as “target instance”. DeepLIFT propagates the predicted variant effect (difference between predicted expression for alternative allele and reference allele) back to input features, resulting in 20,020 attribution scores. We normalized the attribution scores by the total sum and further aggregated by unique epigenomic features (2002 in total).

### Implementing Seal to prioritize causal eQTL variants discovered in fetal and adult brain

We applied the Seal model of 122 brain-region states to eQTL variants from two resources^7,34^. In total, we predicted brain-state-specific expression effects for the following ten sets of fine-mapped eQTL variants (located within 20kb of a gene TSS): 1) 35,193 eQTLs of fetal prefreontal cortex; 2) 28,121 eQTLs of early fetal prefreontal cortex (identified using samples of first-trimester); 3) 17,280 eQTLs of mid fetal prefreontal cortex (identified using samples of second-trimester); 4) 45,920 eQTLs of adult amygdala; 5) 107,249 eQTLs of adult caudate basal ganglia; 6) 150,019 eQTLs of adult cerebellum; 7) 94,997 eQTLs of adult frontal cortex BA9; 8) 70,386 eQTLs of adult hippocampus; 9) 105,757 eQTLs of adult nucleus accumbens basal ganglia; 10) 88,888 eQTLs of adult putamen basal ganglia. We manually mapped the context of each eQTL study to brain-region states of Seal predictions by name. For contexts where exact mapping can not be achieved, we mapped them to the closest subregions (e.g. prefreontal cortex and frontal cortex BA9 were mapped to aggregation of four subregions DFC, MFC, VFC, and OFC; basal ganglia was mapped to striatum). For each eQTL variant, we compared the predicted expression-change direction for the mapped brain-region state with the observed direction from eQTL study. We binned all eQTL variants by eight thresholds of absolute predicted effect (0, 0.02, 0.04, 0.06, 0.08, 0.10, 0.12, 0,14). We computed an accuracy score at each threshold based on the direction comparison for all variants passing the threshold. For contexts that were mapped to multiple brain-region states (prefreontal cortex and frontal cortex BA9), we computed the average accuracy score across all mapped states.

### Partitioning GWAS heritability of brain-associated traits with Seal-predicted variant effects

We downloaded GWAS summary statistics of two brain structural traits^37^ (cortical surface area and cortical mean thickness) and six neuropsychiatric conditions (autism spectrum disorder^63^; attention-deficit hyperactivity disorder^64^; Schizophrenia^65^, bipolar disorder^66^, major depressive disorder^67^, Alzheimer’s disease^43^). We applied the Seal model for 122 brain-region states and 82 cell states of prefrontal cortex to the over 2 million 1000 Genome Project TSS-proximal variants (within 20kb of TSS) analyzed by GWAS studies. To identify specific regulatory programs of each trait, we focused on variants of highly expressed genes in each brain-region or cell state (above 75^th^ percentile). Seal-predicted variant effects for each state were normalized into scores between 0 (minimum) and 1 (maximum). To study the association between Seal-predicted variant effect and trait heritability, we performed GWAS heritability partitioning with the LD-score regression (LDSR) method^36^. We obtained the LDSCORE v2.2 set of known baseline annotations from https://alkesgroup.broadinstitute.org/LDSCORE/. We ran LDSR with Seal-predicted variant effect annotations and a baseline all-ones annotation. We used a conservative enrichment estimator, which is the estimated heritability enrichment score subtracted by one standard error. Benjamini–Hochberg procedure was used to adjust Enrichment P-value for multiple testing on annotations of all traits.

### Analyzing functional burden of SSC de novo mutations with Seal-predicted variant effects

We applied the Seal model of 82 for states of prefrontal cortex to *de novo* mutations discovered in porbands and unaffected siblings from Simons Simplex Cohort^50^. In total, we predicted the cell-state-specific expression effects for 27,926 mutations within 20kb of TSS (out of 127,140 non-repeat-region *de novo* SNVs). We normalized the absolute predicted scores of proband mutations by z-transforming against those of sibling mutations, so that the normalized scores for probands represent deviations from the standardized distribution of sibling mutations (with a mean of 0 and a standard deviation of 1). For functional burden analysis, we focused on the 1,704 mutations (873 proband-specific mutations, 831 sibling-specific mutations) near the 922 SFARI category 1 and 2 genes (defined as genes with presence of recurrent likely gene disrupting mutations). We computed effect size for each cell state as the difference in average normalized scores between proband and sibling mutations. We then performed the one-sided Mann-Whitney U test to compare the normalized scores of proband and sibling mutations by cell state. Benjamini–Hochberg procedure was used to adjust P-value for multiple testing on all cell states.

## Code Availability

The Seal framework code is available at https://github.com/FunctionLab/Seal. The data files required for running Seal framework can be downloaded by following the instructions in the GitHub repository.

## Acknowledgements

We are grateful to the families participating in the SFARI SSC. We thank all the donors for their generosity in tissue donations to the GTEx research project. GTEx datasets used for the analyses described in this manuscript were obtained through dbGaP study accession number phs000424.v10. We thank all lab members in Troyanskaya lab for their constructive feedback. This work was conducted utilizing the computing resources, supported by the Scientific Computing Core, at the Flatiron Institute. This work is supported by funding from the NIH HIGMS no. R01GM071966 and Simons Foundation grant 395506.

## Author contributions

Y.H. and O.G.T. conceived the study. Y.H. designed the study, developed the computational methods and performed the analyses. C.Y.P. and C.L.T. contributed insights, research materials, and analytic tools. Y.H., C.Y.P., C.L.T., and O.G.T. wrote the manuscript.

