## Supplementary Fig. for "Spatiotemporal AI modeling of gene regulation and variant effects in the developing human brain"

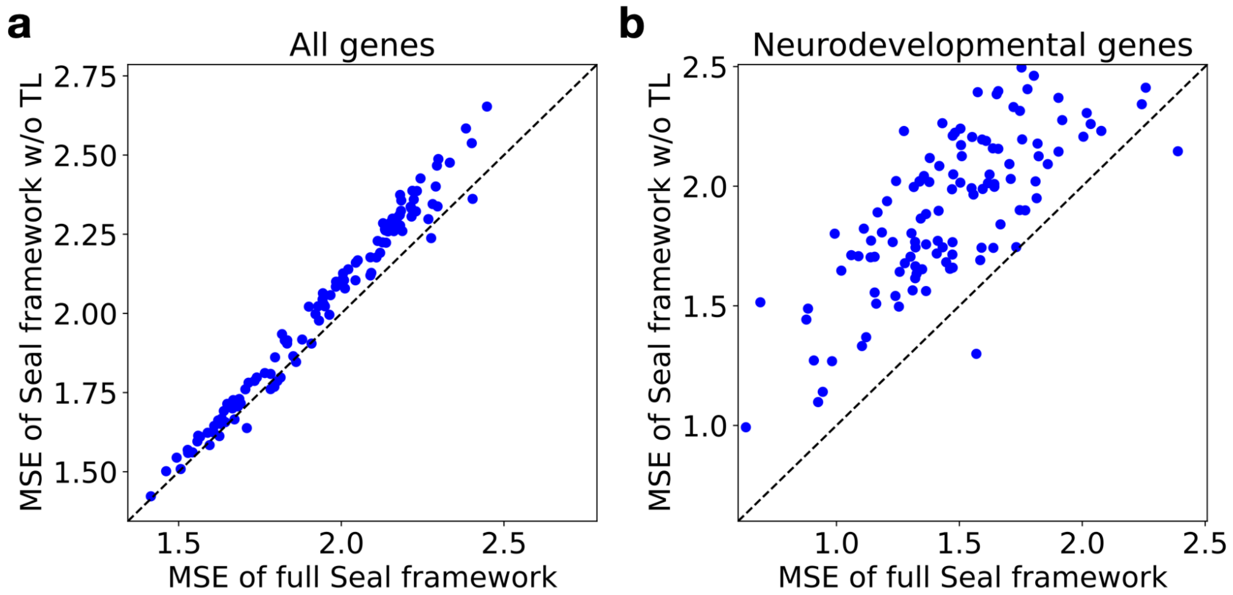


***Supplementary Fig. 1| The transfer learning framework significantly improved the overall model performance of Seal as compared to training with all data in one step***

*The scatter plot displays the comparison of mean squared error (MSE) of Seal (x-axis) compared to the framework without the transfer learning (direct training on all context-specific expression profiles at once, y-axis, (Methods). (**a**) When evaluated on all genes in the testing set, the transfer learning component improved the model performance of Seal, with significantly lower mean squared error (P = 0.01, Wilcoxon signed-rank test). (**b**) The improvement is especially high for the genes involved in neurodevelopment processes (P = 1.1*$10^{-18}$, Wilcoxon signed-rank test).*

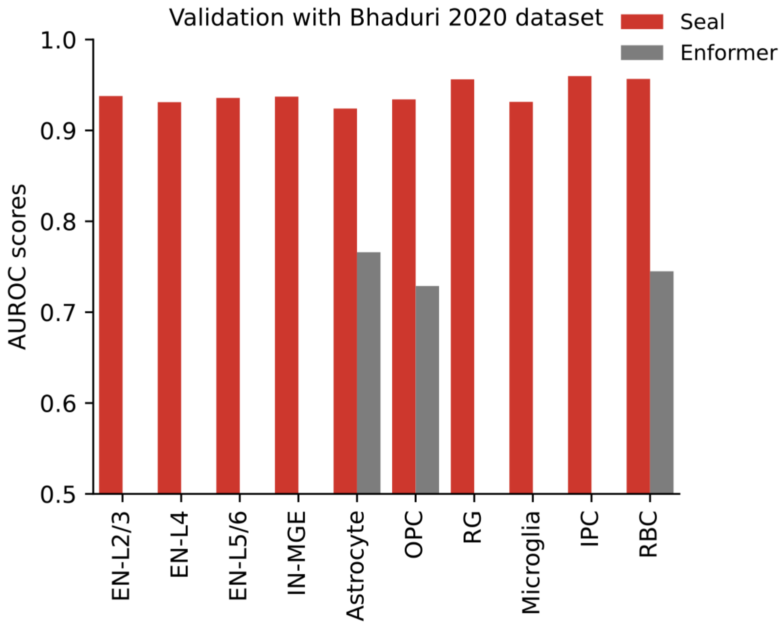


***Supplementary Fig. 2| Seal outperforms Enformer in predicting context-specific gene expression of brain development***

*Sequence-based predictions by Seal and Enformer were benchmarked against an independent single-cell RNA-seq dataset by Bhaduri et al (Validation with the Velmeshev 2019 dataset is shown in Fig. 2). The bar plot shows the computed AUROC of each identified cell state. Seal can make predictions for all the identified cell states while Enformer cannot. Cell states where Enformer cannot make predictions were left blank (no grey bar). (abbreviations, EN: excitatory neuron; IN: interneuron; OPC: oligodendrocyte precursor cell; RG: radial glia; IPC: intermediate progenitor cell; RBC: red blood cell)*

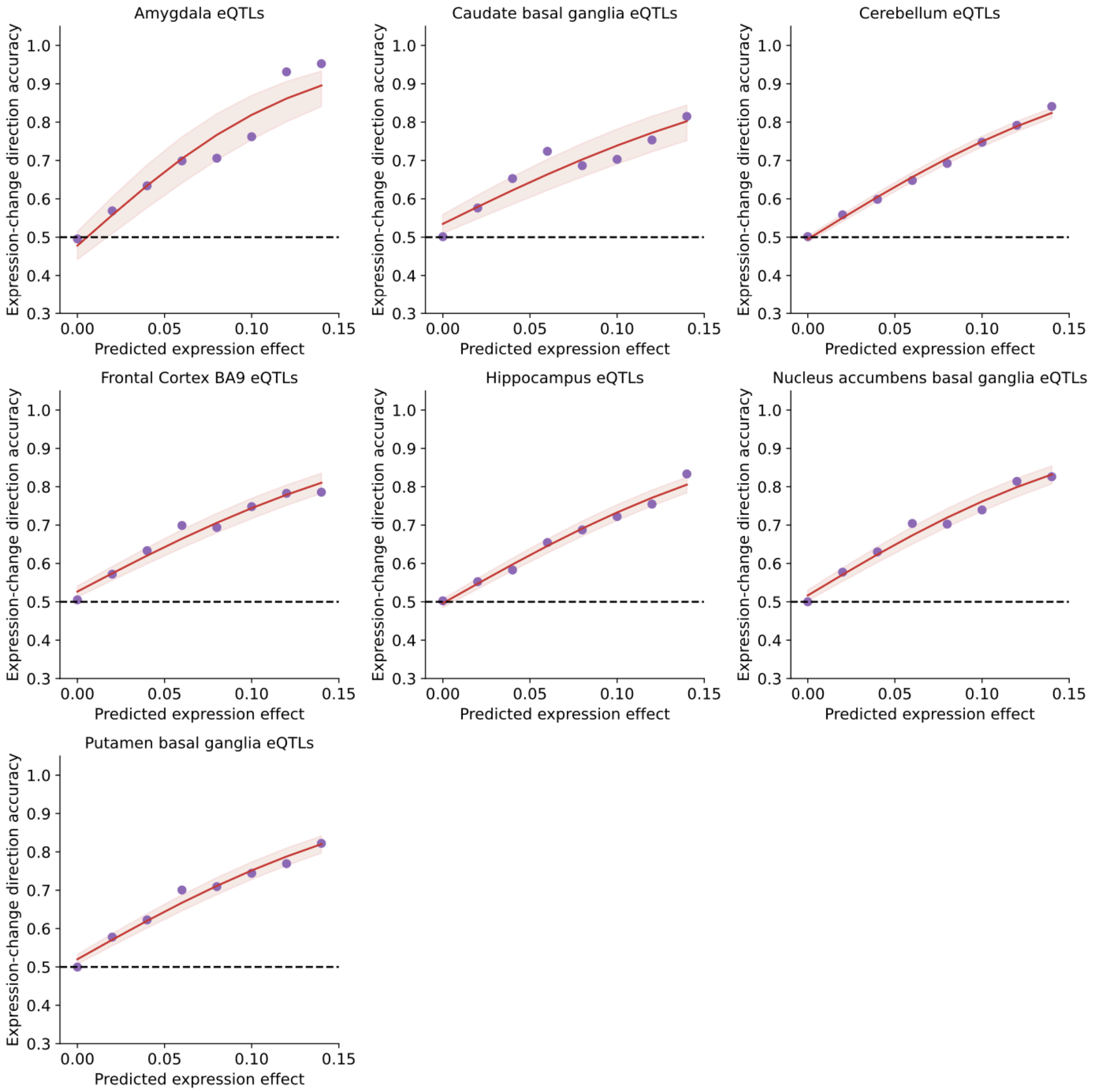


***Supplementary Fig. 3| Seal prioritizes expression-altering eQTL variants for brain development***
*Expression-change direction accuracy increases with the predicted magnitude of variant effect, suggesting eQTLs with strong Seal-predicted effects are more likely to be causal. Each dot in a plot represents the computed accuracy (y axis) for variants above a predicted effect magnitude threshold (x axis, measured by the absolute value of log fold change). Each curve was fitted using the computed accuracy scores at eight different thresholds, with the 95% confidence interval represented by the shaded area. Seal correctly predicted the direction of expression change for top prioritized eQTL variants with high accuracy across all seven adult brain regions (Results for the three sets of eQTL variants discovered from fetal prefrontal cortex can be found in Fig. 3).*

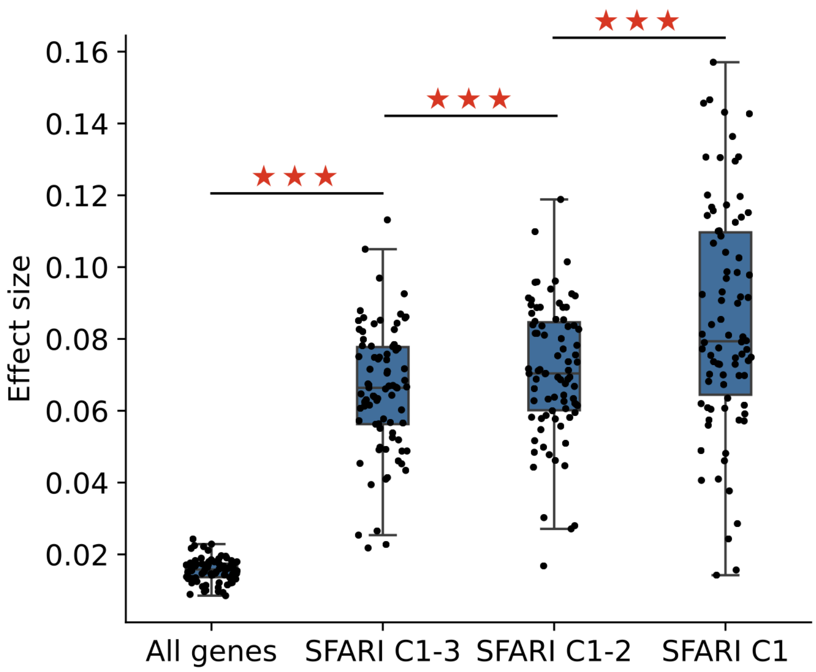


***Supplementary Fig. 4| High functional burden was discovered for proband mutations around high-confidence ASD risk genes***

*Functional burden analysis was conducted to compare the Seal-predicted cell-state-specific mutation effects around ASD risk genes with different levels of confidence. Effect size for a given cell state was computed as the difference in average predicted effects between proband and sibling mutations. The boxplot displays the effect size distribution (y-axis) across cell states for three categories ASD risk genes and all genes (x-axis). SFARI category one genes (SFARI C1) were assigned the highest confidence level by SFARIgene curators, followed by SFARI C2 genes and SFARI C3 genes. The red stars denote the significance level of pairwise comparison by Wilcoxon signed-rank test ($P = 1.5*10^{-4}$ SFARI C1 vs SFARI C1-2; $P = 1.0*10^{-14}$ SFARI C1-2 vs SFARI C1-3; $P = 1.8*10^{-15}$ SFARI C1-3 vs all genes).*
